# The evaluation of the SumiOne^™^ spatial emanators against wild *Anopheles funestus* in experimental huts in Siaya County, western Kenya

**DOI:** 10.64898/2026.09.09.750313

**Authors:** Seline Omondi, Silas Agumba, Mathew Kipsum, Vincent Moshi, Margaret Muchoki, Bernard Abong’o, Brian Polo, Jackline Kosgei, Ashley Hudson, Nicole Achee, Catherine W. Lukhoba, Wolfgang Richard Mukabana, Eric Ochomo

## Abstract

**Background:** Spatial Emanators (SEs) are a promising tool for complementing existing malaria vector control interventions. In 2025, the World Health Organization (WHO) issued a conditional recommendation for SEs, and two transfluthrin-based products were added to the WHO Prequalification List. As SEs are scaled up, additional products are needed to ensure reliable supply, competitive pricing, and continued innovation.

**Methods:** This study evaluated the efficacy of SumiOne™ SE, containing 10% metofluthrin, in experimental huts in Siaya County, western Kenya. SumiOne™ SE efficacy was evaluated in six experimental huts, three intervention and three control, using human landing catches (HLCs) and aspiration. Collections were conducted in two 12-night phases, alternating between HLC and aspiration every six nights. Outcomes included reductions in human landing rates, density, exophily and blood feeding inhibition of *Anopheles* mosquitoes.

**Results:** SumiOne™ SE significantly reduced indoor densities of *An. funestus* with 63% protective effect (IRR = 0.37, 95% CI: 0.20-0.69, p = 0.0087). Overall *An. funestus* density (indoor and outdoor combined) was reduced by 58% (IRR=0.42, 95% CI: 0.22–0.79; p=0.0222). The rate of exophily of *An. funestus* exiting was 34% (95% CI: 27-42) in the intervention huts compared to 2% (95% CI: 1-4) in the control huts (p< 0.0001). Blood feeding inhibition for *An. funestus* was 46% with 44% (95% CI: 36-51) of the collected mosquitoes being blood fed in the intervention huts compared to 81% (95% CI: 76-86) blood fed in the control huts, (p<0.0001).

**Conclusion:** These findings support SumiOne™ SE’s potential as a complementary vector control tool, alongside existing core interventions such as Insecticide Treated Nets and Indoor Residual Spraying, particularly in contexts where additional protection is needed.

## Introduction

Despite extensive efforts to control malaria-transmitting mosquitoes through widespread distribution of insecticide-treated nets (ITNs) and indoor residual spraying (IRS), malaria remains a significant public health challenge, particularly in sub-Saharan Africa (SSA) (1). In 2024, there were 282 million cases of malaria and 610,000 deaths (1). This is mainly attributed to the high efficiency of vectors, such as *Anopheles gambiae* and *Anopheles funestus*, in transmitting *Plasmodium falciparum* parasites, which accounts for the vast majority of malaria morbidity and mortality in sub-Saharan Africa (2). Compounding the problem is the rapid spread and high intensity of insecticide resistance among these mosquito populations, which undermines the effectiveness of key active ingredients in most of the ITNs (3). Because pyrethroid insecticides remain the primary active ingredient class used in most ITNs, resistance directly compromises both personal and community-level protection, creating a substantial gap in current malaria control strategies related to resistance.

In parallel with physiological resistance, malaria vectors also exhibit behavioral adaptations that further limit the effectiveness of ITNs and IRS. Recent studies have reported shifts in mosquito biting behavior, with *Anopheles* species increasingly found host-seeking during the day, both indoors and outdoors. For instance, Omondi et al.,2023(4) reported late-morning biting in *An. funestus* in western Kenya, with significant activity occurring between 06:00 and 09:00 hours, well after the typical overnight peak. This shift in biting behavior may reduce the effectiveness of malaria control interventions, such as ITNs and IRS, which are designed to target nocturnal, indoor-biting mosquitoes. Similarly, Odero et al., 2024(5) reported that *An. gambiae s*.*s*. in western Kenya actively bite outdoors between 5:00 AM and 7:00 AM, increasing transmission risk for individuals engaged in early-morning activities, including preparing for school, cleaning compounds, working on farms, and milking cows. Similarly, studies from other African settings have reported extended biting activity among malaria vectors, including outside the traditional sleeping hours. In Tanzania a 24-hour biting pattern was documented for *An. arabiensis* (6). Likewise, in Bangui, Central African Republic, approximately 20–30% of indoor biting by *An. gambiae, An. coluzzii, An. funestus*, and *An. pharoensis* occurred during daytime hours (7). These gaps in protection call for complementary interventions that target mosquitoes biting outside the protection provided by current indoor-based interventions.

Spatial Emanators (SEs) are volatile chemical compounds that reduce human-vector contact by interfering with the host-seeking behavior of haematophagous insect such as mosquitoes (8). Commonly used SE active ingredients include pyrethroid-based compounds such as transfluthrin and metofluthrin, which are characterized by high vapor pressure and rapid volatilization at ambient temperatures (9). Spatial emanators have potential advantages over existing control measures, such as ITNs and IRS, in that they are easier to use and deploy, require limited behavioral modification of household residents, and are effective when individuals are not protected by indoor vector control tools(10-12).

In 2025, the WHO issued a conditional policy recommendation for SEs, and two SE products with transfluthrin as the active ingredient (Mosquito Shield^™^ and Guardian™) have been included on the WHO Prequalification List (1). Given the WHO conditional recommendation and subsequent scale-up of SEs, multiple SE products are needed to ensure consistent supply chains and competitive pricing while spurring innovation for novel SE products. This study evaluated the efficacy of the SumiOne™, an SE product with metofluthrin as its active ingredient, against free-flying pyrethroid-resistant *Anopheles* mosquitoes inside and outside experimental huts based on entomological indicators.

## Methodology

### Study site

The study was conducted in Ruambwa experimental huts, which are situated within the Bunyala Rice Irrigation Scheme. These experimental huts are located near rice paddies in the Alego-Usonga sub-county, Siaya County, western Kenya. The availability of water, combined with high temperatures year-round, creates an ideal environment for the proliferation of *Anopheles* malaria vectors throughout the year. This area is characterized by high populations of both *An. funestus* and *An. arabiensis*, with very high levels of pyrethroid resistance(13).

There are 17 experimental huts in the Ruambwa, spaced 20 m apart, with each hut measuring 6 m in length, 3 m in width, and 2 m in height. The experimental huts design resembles the ones described in Agumba et al., 2024 (10) and Kipsum et al., 2026 (14) with the only addition that they feature a veranda measuring 3.5 m in width and 6 m in length, providing shade for volunteers during outdoor HLC collections conducted in the day.

### Study design and intervention

SumiOne™ SE is formulated as a plastic mesh incorporated with 10% metofluthrin 10% as its active ingredient (Sumitomo Chemical Company Ltd., Chuo-ku, Tokyo, Japan). Four SumiOne™ SEs were hung per hut, i.e., one device per approximately 5 m^2^, using a piece of string tied to a hook to suspend the emanator from the ceiling of the hut. The products were placed 1.52m apart and suspended at a height of 1.5 m from the floor.

Six huts were selected and randomly allocated to either SumiOne™ SE treatment or controls, (3 huts per arm). The evaluation consisted of 24 collection nights, split into two phases of 12 days each. Each phase consisted of two cycles of six collection days, alternating between HLC and aspiration collections following overnight sleepers every six days.

### Baseline survey

Baseline mosquito collection was conducted using HLC for 18 h (18:00 to 12:00) for five consecutive days in January 2025 prior to the intervention. The mosquito collections were then conducted from January to March 2025.

### Randomisation

Randomisation of sleepers was carried out by assigning each sleeper a number corresponding to one of the six huts. For each 6-day cycle of collections, sleepers were rotated among the huts such that each sleeper occupied each hut for one night per cycle. Allocation was conducted using a simple lottery method.

Randomisation of huts to treatment and control arms was performed to balance mosquito densities observed during baseline collections, resulting in huts 1, 3 and 5 assigned to the intervention arm, and huts 2, 4, and 6 assigned to the control arm.

### Study participants

The study was conducted in accordance with WHO guidelines for evaluation of spatial emanator products (15). Adult humans (aged 18–45 years) were recruited as paid mosquito collectors. Written informed consent was obtained from all mosquito collectors who participated in the study. Collectors were provided with doxycycline as malaria prophylaxis for the duration of the study. All collectors were also tested for malaria during recruitment and at any time they became ill during the study. Those found positive with malaria were treated using artemether-lumefantrine (Coartem®) according to the Kenya National Guidelines(16).

### Human landing catches

Human landing catches (HLC) were conducted for 18 h (18:00 h to 12:00 h), both indoors and outdoors in the experimental huts. A total of 36 study participants were organized into groups of six (three inside and three outside on the verandas) at the start of the experiment and remained in the same group for the duration of the study. Each night’s collection period was divided into three 6-h shifts: 18:00–00:00 h, 00:00–06:00 h, and 06:00–12:00 h (the last two shifts happening the next day).

The 36 study participants rotated among the huts nightly, ensuring that each group collected mosquitoes in each hut. The three participants who rotated as a group alternated among the three collection shifts in a staggered manner.

During the mosquito collection process, collectors sat on chairs wearing pairs of short trousers and long-sleeved shirts and used a mouth aspirator (Model 412, John W. Hock Company, Gainesville, Florida, USA) to collect mosquitoes landing on their lower legs. Landed mosquitoes were aspirated before biting. Collected mosquitoes were placed in clean paper cups, which were changed hourly and by location. Mosquitoes were provided with access to a 10% sugar solution and transported in a cooler box to the field laboratory for processing. In the laboratory, the *Anopheles* mosquitoes were separated by species, sex and abdominal status (blood-fed, unfed, gravid or half-gravid), and the number collected per hour was recorded. Morphological identification was performed using standard taxonomic keys [20] to differentiate *An. funestus* sensu lato and *An. gambiae* sensu lato and other secondary malaria vectors. A subset of the *An. funestus* sensu lato and *An. gambiae* sensu lato were identified to species using Polymerase Chain Reaction (17, 18).

### Blood-feeding evaluations

Blood-feeding evaluations were carried out for 11 h (20:30 h to 06:30 h) per night during the study period. Six study participants rotated among the six huts each night, sleeping under untreated bed nets from 20:30 h until 06:30 h. To simulate wear and tear following WHO guidelines, each net was deliberately holed with six 4 cm × 4 cm holes (two on each long side and one on each short side) [21].

Mosquitoes were collected each morning between 06:30 h and 07:30 h using mouth aspirators from different locations, including nets, under the bed, roof, floor, walls and exit traps, and transported to the field laboratory. Mosquitoes were classified as dead or alive and by abdominal status (unfed, fed, gravid or half-gravid). Knockdown and mortality were recorded one-hour post-collection. Live mosquitoes were maintained at 27 ± 2 °C with access to 10% sugar solution for up to 24 h to assess delayed mortality. Female mosquitoes were identified to species level using morphological keys [20].

### Statistical analysis

Vector species abundance was assessed using descriptive statistics (means, proportions and 95% confidence intervals [CI]). Generalized linear mixed models (GLMM) using Template Model Builder (package glmmTMB) were fitted using a negative binomial distribution for analysis of mosquito numbers at different collection locations (indoors or outdoors). Models were adjusted for repeated measures using the collection hut ID as a random effect. Model coefficients were exponentiated to determine the incidence risk ratios (IRR) and their 95% confidence intervals. The statistical significance level was set at α = 0.05. The protective efficacy for each experiment was calculated as (1 − IRR) * 100, where IRR was the incidence rate ratio in the SumiOne™ spatial emanators group compared to the control. All data analyses were performed using R statistical software version 4.1.2.

Deterrence, blood-feeding inhibition, and personal protection were calculated as follows:

- Exophily – proportion of mosquito in exit traps
- Deterrence (%) = ((Nc − Nt) / Nc) × 100
- Blood-feeding inhibition (%) = ((PBc − PBt) / PBc) × 100
- Personal protection (%) = ((Bc − Bt) / Bc) × 100

Where:

Nc = number of mosquitoes in the control huts exit traps

Nt = number of mosquitoes in the intervention huts exit traps

PBc = proportion of blood-fed mosquitoes in control huts

PBt = number of blood-fed mosquitoes in intervention huts

Bc = number of blood-fed mosquitoes in control huts

Bt = number of blood-fed mosquitoes in intervention huts

## Results

### Baseline collections

A total of 603 female *Anopheles* mosquitoes were collected using HLC during the five-day baseline survey. Morphological identification showed that most were *An. funestus* (n=512, 84.9%). *An. gambiae* (n=72, 11.9%) was also recovered. Other species that were collected included *An. Coustani* (1), *An. Pharoensis* (14), and *An. Ziemanni (4)*. PCR-based species identification indicated that all tested specimens within the *An. funestus* group were *An. funestus sensu stricto* (100%). Among the *An. gambiae sensu lato* complex, 65% were *An. arabiensis*, while the remaining 35% were *An. gambiae sensu stricto*. For location, 383 *An. funestus* were collected indoors while 129 were collected outdoors. For *An. gambiae*, 35 were collected indoors while 37 were collected outdoors (Table 1). There was no difference in the mean biting activity among the six huts (all pairwise p-values > 0.05), and the huts were balanced for the intervention and control arms.

**Table 1.** Female *Anopheles* and *Culicine* species composition during baseline collections.

| Species | Indoor |  | Outdoor |  |
| --- | --- | --- | --- | --- |
|  | n (%) | Mean/hut/night | n (%) | Mean/hut/night |
| *An. funestus* | 383 (15.8) | 12.8 | 129 (8.3) | 4.3 |
| *An. gambiae* | 35 (1.4) | 1.2 | 37 (2.4) | 1.2 |
| *An. coustani* | 0 | 0 | 1 (0.1) | 0.03 |
| *An. pharoensis* | 2 (0.1) | 0.1 | 12 (0.8) | 0.4 |
| *An. ziemanni* | 2 (0.1) | 0.1 | 2 (0.1) | 0.1 |
| *Culicines* | 1996 (82.5) | 66.5 | 1364 (88.3) | 45.5 |

**Table 2.** Summary statistics of SumiOne™ in reducing human landings of wild *Anopheles* mosquitoes.

| Category | Location | Arm | Mean/hour<br>(se) | IRR<br>(95% CI) | Mean difference<br>(95% CI) | p.value |
| --- | --- | --- | --- | --- | --- | --- |
| *An.*<br>*funestus* | Indoor | Intervention | 0.21 (0.05) | 0.37 (0.20-0.69) | 0.34 (0.09-0.60) | **0.0087** |
|  |  | Control | 0.55 (0.13) | Ref | Ref |  |
|  | Outdoor | Intervention | 0.18 (0.05) | 0.56 (0.31-1.00) | 0.14 (-0.01-0.30) | 0.0702 |
|  |  | Control | 0.33 (0.09) | Ref | Ref |  |
|  | Overall | Intervention | 0.14 (0.04) | 0.42 (0.22-0.79) | 0.20 (0.03-0.37) | **0.0222** |
|  |  | Control | 0.34 (0.09) | Ref | Ref |  |
| *An.*<br>*gambiae* | Indoor | Intervention | 0.10 (0.05) | 0.83 (0.53-1.31) | 0.02 (-0.03-0.07) | 0.4572 |
|  |  | Control | 0.12 (0.06) | Ref | Ref |  |
|  | Outdoor | Intervention | 0.41 (0.27) | 1.02 (0.66-1.59) | -0.01 (-0.19-0.17) | 0.9282 |
|  |  | Control | 0.40 (0.25) | Ref | Ref |  |
|  | Overall | Intervention | 0.11 (0.14) | 0.90 (0.59-1.35) | 0.01 (-0.04-0.07) | 0.6546 |
|  |  | Control | 0.12 (0.16) | Ref | Ref |  |
| Female<br>*Anopheles* | Indoor | Intervention | 0.237 (0.051) | 0.41 (0.234-0.717) | 0.34 (0.100-0.583) | **0.0057** |
|  |  | Control | 0.578 (0.115) | Ref | Ref |  |
|  | Outdoor | Intervention | 0.181 (0.035) | 0.67 (0.410-1.108) | 0.09 (-0.025-0.2) | 0.1286 |
|  |  | Control | 0.268 (0.049) | Ref | Ref |  |
|  | Overall | Intervention | 0.196 (0.041) | 0.49 (0.282-0.854) | 0.20 (0.031-0.375) | **0.0208** |
|  |  | Control | 0.398 (0.079) | Ref | Ref |  |

### SumiOne™ SE evaluation

#### Human landing catches

A total of 1486 female *Anopheles* mosquitoes were collected by HLC over 12 days post-intervention deployment. The intervention significantly reduced the indoor and outdoor densities of *An. funestus* with 63% protective effect indoors (IRR = 0.37, 95% CI: 0.20-0.69, p = 0.0087) while there was a 44% reduction in outdoor *An. funestus* although the effect was not statistically significant (IRR = 0.56, 95% CI: 0.31-1.00, p = 0.0702). The intervention significantly reduced the overall (indoor + outdoor) *An. funestus* density with an IRR of 0.42 (95% CI: 0.22-0.79, p = 0.0222), corresponding to a 58% protective effect. For *An. gambiae*, there were no statistically significant differences compared to the control huts either indoors or outdoors. The overall IRR for *An. gambiae* was 0.90 (95% CI: 0.59-1.35, p = 0.6546). For overall female *Anopheles* mosquitoes, SumiOne™ SE significantly reduced indoor landing rates, with an IRR of 0.41 (95% CI: 0.234–0.717, p = 0.0057), corresponding to a 59% protective efficacy compared to control huts. When indoor and outdoor data were combined, an overall 51% reduction in landing rates was observed (IRR = 0.49, 95% CI: 0.282–0.854, p = 0.0208) (Table 1).

#### Evaluation of blood-feeding inhibition

A total of 556 female *Anopheles* mosquitoes were collected during the evaluation of the impact of SumiOne™ SE’s on blood feeding inhibition. Of these, 255 *An. funestus* were collected in the control huts, compared with 174 collected from the intervention huts. In contrast, more *An. gambiae* were collected from the intervention huts than from the control huts (86 vs 41). Overall, SumiOne™ SE provided 54% personal protection. Species-specific analysis indicated that SumiOne™ SE provided 63% personal protection against *An. funestus* while there was no evidence of personal protection observed against *An. gambiae*. However, significantly fewer *Anopheles* (*An. funestus* + *An. gambiae*) were blood-fed in the intervention huts (p<0.0001). Blood feeding inhibition for *An. funestus* was 46% with 44% (95% CI: 36%-51%) of the collected mosquitoes being blood fed in the intervention huts compared to 81% (95% CI: 76-86) blood fed in the control huts. For *An. gambiae*, blood feeding inhibition was 32% with 51% (95% CI: 35%-67%) blood fed in the intervention huts compared to 35% (95% CI: 25-46) in the control huts, though the difference was not statistically significant (p=0.2821).

For *Anopheles* mosquitoes exiting the huts, there was a significantly higher exit rate of 38% (95% CI: 32%-44%) in the intervention huts compared to 7% (95% CI: 4%-10%) in the control huts (p< 0.0001). For *An. funestus*, the rate of exophily was 34% (95% CI: 27%-42%) in the intervention huts compared to 2% (95% CI: 1%-4%) in the control huts (p< 0.0001).

Observed 24-hour mortality was significantly higher in control huts compared to intervention huts, 20% vs 12% (p = 0.0281) (Table 3).

**Table 3.** Effect of SumiOne™ Spatial emanator on *Anopheles funestus* and *Anopheles gambiae* exophily (exit rate), blood feeding inhibition, and mortality.

| Species | Arm | n | %<br>Deterrence | P-value | %<br>Exophily | 95% CIs | % Blood<br>feeding | 95% CIs | P-value | % Blood<br>feeding<br>inhibition | %<br>Personal<br>protection | % 24h<br>mortality |
| --- | --- | --- | --- | --- | --- | --- | --- | --- | --- | --- | --- | --- |
| *An. funestus* | Intervention | 174 | 32 | <0.0001 | 34 | 27-42 | 44 | 36-51 | <0.0001 | 46 | 63 | 10 |
|  | Control | 255 | - | Ref | 2 | 1-4 | 81 | 76-86 | Ref | - | - | 18 |
| *An. gambiae* | Intervention | 86 | -110 | 0.2824 | 47 | 36-58 | 51 | 35-67 | 0.2821 | 32 | -43 | 16 |
|  | Control | 41 | - | Ref | 39 | 25-55 | 35 | 25-46 | Ref | - | - | 27 |
| Female<br>*Anopheles* | Intervention | 260 | 12 | <0.0001 | 38 | 32-44 | 41 | 35-47 | <0.0001 | 47 | 54 | 12 |
|  | Control | 296 | - | Ref | 7 | 4-10 | 77 | 72-82 | Ref | - | - | 20 |

## Discussion

This study evaluated the efficacy of SumiOne™ SE, a metofluthrin-based spatial emanator, against free-flying pyrethroid-resistant *Anopheles* mosquitoes under experimental hut conditions in Siaya County, western Kenya. The findings demonstrate that SumiOne™ SE substantially reduced mosquito–human contact, including significant reductions in landing rates and blood-feeding, as well as increased exiting behaviour. These findings are consistent with the established mode of action of spatial repellents [13] and support the potential of SumiOne™ SE as a complementary vector control tool in settings where residual transmission persists despite high coverage of conventional interventions.

Species-specific analyses showed that the reduction in mosquito density was primarily driven by effects on *An. funestus*. This is particularly important given the predominant role of *An. funestus* in malaria transmission in western Kenya and its well-documented high levels of pyrethroid resistance. The strong efficacy against this species suggests that spatial repellents may be particularly valuable in settings where conventional insecticide-based tools may be compromised by resistance. The behavioral mode of action of metofluthrin, which does not rely on mosquito mortality, may explain why efficacy was maintained despite high resistance intensity. The efficacy observed for *An. funestus* is consistent with previous experimental hut evaluations of metofluthrin- and transfluthrin-based spatial repellents [9, 14, 15].

The lack of a significant effect on *An. gambiae* s.l. may be due to several factors. During the study period, *An. gambiae* density at the site was substantially lower than that of *An. funestus*, reducing statistical power to detect a moderate effect. Additionally, the predominant sibling species within *An. gambiae* s.l. at this site is *An. arabiensis* [12], a species with comparatively exophilic and exophagic behaviour that may result in reduced indoor exposure to airborne metofluthrin concentrations. It is also possible that the null result reflects a genuine difference in sensitivity between species to metofluthrin at the doses generated by this product formulation, or differences in flight behaviour leading to differential exposure. Future studies with larger sample sizes and species-specific dose-response data would be needed to resolve this question definitively.

SumiOne™ SE also substantially inhibited blood feeding among collected *Anopheles* mosquitoes. Blood-feeding inhibition directly reduces the probability of parasite transmission, thereby lowering vectorial capacity even in the absence of high mosquito mortality. The observed blood-feeding inhibition of *An. funestus* suggests that SumiOne™ SE has the potential to reduce human–vector contact by decreasing the proportion that successfully obtain a blood meal. Although personal protection was also observed, this metric is influenced not only by blood-feeding inhibition but also by differences in mosquito densities between intervention and control huts. Consequently, the two measures should be interpreted as complementary rather than interchangeable indicators of intervention performance. Blood-feeding inhibition is a critical entomological endpoint because it directly reduces vectorial capacity [16]. This level of protection may be substantial, particularly in areas with low ITN use or biting outside the times when people are protected by ITNs. The observed efficacy was achieved against a highly pyrethroid-resistant vector population, supporting the hypothesis that spatial repellency acts through non-toxic behavioural mechanisms unlikely to be directly undermined by the metabolic or target-site resistance mechanisms that compromise pyrethroid-based ITNs and IRS [9]. The reduction in blood-feeding observed here aligns with previous findings from spatial repellent evaluations in East Africa and South-East Asia [10, 14].

The marked increase in mosquito exiting behaviour in intervention huts further supports the behavioural mode of action of metofluthrin. The exit rate of all *Anopheles* mosquitoes in intervention huts was high compared to that in control huts, with *An. funestus* showing an exophily rate of 34% versus 2% in controls. This is consistent with findings from experimental hut trials of other volatile pyrethroids, where transfluthrin and metofluthrin induced exiting rates of approximately 38% and 30%, respectively, in *An. gambiae* [17, 18]. Similar patterns have been reported for transfluthrin-based spatial emanators, including SC Johnson Guardian™ and Mosquito Shield™ [14, 18]. The observation that blood-feeding rates were substantially combined with high exophily rates indicates that mosquitoes are deterred before obtaining a bloodmeal. In this study, exiting mosquitoes were captured in traps. However, in theory, exiting mosquitoes could be diverted to neighboring houses, particularly those that do not have SEs. A trial of spatial emanators (Mosquito Shield) in western Kenya did not find evidence of diversion based on entomological or epidemiological outcomes(19). However, it is possible that mosquitoes respond differently to transfluthrin versus metofluthrin and further studies are needed to estimate the risk of transmission by mosquitoes diverted from homes that have SEs.

The limited insecticidal activity observed in the present study contrasts with some evaluations of metofluthrin- and transfluthrin-based SEs that have reported appreciable knockdown and mortality in addition to behavioral effects. For example, metofluthrin vaporized using an LPG-powered emanator induced high levels of knockdown and mortality against *An. funestus* in experimental huts in western Kenya, while a transfluthrin-based passive emanator evaluated in Benin also produced measurable mortality against pyrethroid-resistant *An. gambiae* s.l. (10, 20). Collectively, these findings suggest that mortality is not a universal outcome of SEs evaluation and may depend on factors such as product formulation, active ingredient release rate, exposure duration, mosquito species, insecticide susceptibility, and environmental conditions. Further studies are therefore needed to better characterize the mechanisms by which different SE formulations influence mosquito behavior and survival under operational conditions.

The study has important operational strengths. It was conducted at a well-characterized site with documented very high pyrethroid resistance levels [12], against free-flying wild vector populations, and a controlled experimental-hut study with daily sleeper rotation to control for collector attractiveness. The evaluation measured multiple complementary endpoints — landing inhibition, blood-feeding inhibition, exophily, and mortality — allowing a comprehensive characterization of the product’s entomological mode of action. The use of both HLC and aspiration collection methods provided triangulation across different aspects of mosquito behavior, and the testing conditions were in line with WHO evaluation guidelines for spatial repellent products.

Nonetheless, several important limitations must be acknowledged. First, the evaluation was conducted in experimental huts rather than real dwellings, and efficacy under programmatic conditions with variable product use may differ. Second, the use of only three huts per study arm limited statistical power, particularly for species-specific analyses, while the lack of hut rotation did not control for potential hut effects. Third, the study had a short evaluation period, confined to a single site and season and environmental conditions such as ambient temperature and relative humidity were not measured. These factors may influence metofluthrin volatilisation and dispersion and, consequently, its efficacy. Efficacy may also vary with seasonal changes in vector density and differences in house structure. Fourth, no epidemiological endpoints were measured, and translation of entomological effects into reduced malaria incidence remains to be established. Lastly, the potential for community-level diversion of mosquito biting to unprotected individuals was not assessed; this is a critical consideration for SEs, and monitoring programs should be designed to address this as SEs are scaled up.

These findings are timely given the 2025 WHO conditional recommendation for spatial SEs as a supplementary malaria vector control tool [1, 13]. Both currently WHO-prequalified products — Mosquito Shield™ and Guardian™ — contain transfluthrin as the active ingredient. Ensuring that alternative, metofluthrin-based products such as SumiOne™ are rigorously evaluated and available in the market is important for maintaining supply chain resilience, enabling competitive pricing, and mitigating over-reliance on a single active ingredient.

In conclusion, SumiOne™ SE significantly reduced *An. funestus* landing rates, inhibited blood feeding, and increased mosquito exiting behavior in a high-pyrethroid-resistance setting in western Kenya. Mosquito mortality was not affected, suggesting this product acted primarily through behavioral deterrence. These findings support its potential as a complementary vector control tool alongside ITNs and IRS in settings where residual transmission persists. As spatial emanators are scaled up for the prevention of malaria and other vector-borne diseases, future monitoring and evaluation activities should include assessment of community-level diversion, investigation of differential species sensitivity, evaluation of user acceptability and consistent product use under programmatic conditions, and post-deployment safety monitoring and assessment data on household metofluthrin exposure in sub-Saharan African contexts.

## Ethical approval

This study was reviewed and approved by the KEMRI Scientific and Ethics Review Unit, SERU 4899. Written informed consent was obtained from all mosquito collectors who participated in the study.

## Acknowledgment

We would like to thank the study volunteers who participated in the experimental hut trials. We would also like to thank the Asembo laboratory team for their support in supervising the HLC volunteers and processing the samples.

## Funding

This project was made possible thanks to Unitaid’s funding and support. Unitaid saves lives by making new health products available and affordable for people in low- and middle-income countries. Unitaid works with partners to identify innovative treatments, tests and tools, help tackle the market barriers holding them back, and get them to the people who need them most – fast. Since it was created in 2006, Unitaid has unlocked access to more than 100 groundbreaking health products to help address the world’s greatest health challenges, including HIV, TB, and malaria; women’s and children’s health; and pandemic prevention, preparedness and response. Every year, these products benefit more than 300 million people. Unitaid is a hosted partnership of the World Health Organization

## Notes

### Competing Interest Statement

The authors have declared no competing interest.

